# VEGF-C expression in TAMs rewires the metastatic fate of breast cancer cells

**DOI:** 10.1101/2022.01.26.468593

**Authors:** Kaveri Banerjee, Thomas Kerzel, Tove Bekkhus, Sabrina de Souza Ferreira, Tatjana Wallmann, Majken Wallerius, Laura-Sophie Landwehr, Dennis Alexander Agardy, Nele Schauer, Anna Malmerfeldt, Jonas Bergh, Margarita Bartish, Johan Hartman, Arne Östman, Mario Leonardo Squadrito, Charlotte Rolny

**Author notes:** Equal Contribution (Thomas Kerzel, Tove Bekkhus, Sabrina de Souza Ferreira, Tatjana Wallmann). Corresponding authors: Charlotte Rolny, Karolinska Institutet, Department of Oncology-Pathology, SciLifeLab, Solna 171 76 Stockholm; Mario Leonardo Squadrito^2^ San Raffaele Telethon Institute for Gene Therapy (TIGET), Universitá Vita-Salute San Raffaele, Milan, Italy.

## Abstract

Expression of pro-lymphangiogenic vascular endothelial growth factor C (VEGF-C) in primary tumors correlates with the occurrence of proximal lymph node metastasis in most solid cancer types. However, the role of VEGF-C in regulating tumor cell dissemination to distant organs is currently unclear. Perivascular tumor-associated macrophages (TAMs) are key regulators of hematogenous cancer cell spreading, forming tumor microenvironment of metastasis (TMEM) doorways for breast cancer cells to intravasate tumor blood vessels and fuel distant metastases. Using an experimental breast cancer (BC) model, we show here that TAMs expressing VEGF-C decrease cancer cell dissemination to the lung while enhancing lymph node metastasis. These TAMs express podoplanin and associate with normalized tumor blood vessels expressing VEGFR3. Further clinical data reveal that VEGF-C^+^ TAMs correlate inversely with malignant grade and with the occurrence of TMEM complexes in a cohort of BC patients. Thus, our study displays an apparently paradoxical role of VEGF-C expressing TAMs in redirecting cancer cells to preferentially disseminate to the lymph nodes, at least in part, by normalizing tumor blood vessels and promoting lymphangiogenesis.

## Introduction

Breast cancer (BC) is one of the most prevalent cancers worldwide. ^1^ While the prognosis for non-macrometastatic BC is generally good due to availability of efficient adjuvant therapies, including targeted and endocrine therapy, the prognosis for metastatic BC or triple-negative breast cancer (TNBC) is worse partly due to resistance to therapy. New treatment strategies and prognostic biomarkers for these patients are therefore required.

VEGF-C promotes tumor lymphangiogenesis via its receptor VEGFR3, facilitating cancer cell dissemination to the sentinel lymph nodes. ^2^ Even though the occurrence of metastases in tumor-draining lymph nodes is associated with poor clinical outcomes for many solid cancers, including BC, ^3^ it is still unclear to what extent metastatic seeding to lymph nodes further fuels pulmonary cancer cell colonization. ^4,5^ Experimental models of melanoma and mammary carcinoma suggest that a fraction of cancer cells populating lymph nodes do invade lymph node blood vessels, enter the blood stream circulation and colonize the lung. ^6,7^ On the other hand, by performing genetic analyses of clonal evolution between the primary tumor, lymph node metastases, and distant metastases of BC patients, we showed that cancer cells residing at distant sites stem from the primary tumor and not from the sentinel lymph nodes, ^8^ suggesting that BC cells disseminate mainly via the hematogenous route to colonize distant organs.

Abnormal expression of VEGF-A results in atypical tumor vessels that are usually irregular, disorganized, permeable, and leaky, thus facilitating cancer cell intravasation into blood vessels. ^9^ In contrast, normalization of the tumor vasculature is characterized by enhanced vessel perfusion, reduced tumor hypoxia and increased pericyte coating, resulting in hampering of distant metastasis. ^10^ Perivascular tumor associated-macrophages (TAMs) are a major source of VEGF-A, and act as “doorways” for cancer cells to intravasate blood vessels. ^11^ Indeed, TAMs play a key role in micro-anatomical structures termed tumor microenvironment of metastasis (TMEM), which are associated with increased occurrence of metastasis and are composed of a cancer cell (expressing the actin-regulatory protein mammalian-enabled, MENA), a perivascular TAM and an endothelial cell. ^12^ The density of TMEM assemblies has been shown to correlate with the occurrence of distant metastases in experimental mammary tumor models ^11^ and BC patients. ^12–14^ For this reason, reprograming TAMs from a pro-angiogenic to an angiostatic state ^15^ or impairing their association to blood vessels ^16^ may be promising for anti-metastatic interventions. ^15,16^

Building on the fact that TAMs are key players in BC metastatic dissemination, we investigated which TAM-derived factors could regulate the route of metastatic dissemination. We found that TAMs derived from a mammary tumor model preferentially disseminating via the lymphatic route expressed higher levels of VEGF-C as compared to TAMs isolated from a tumor model that preferentially disseminated via the hematogenous route. In turn, forced overexpression of VEGF-C in TAMs negatively regulate number of lung metastases by switching the route of metastatic dissemination from hematogenous to lymphatic. The mechanism whereby TAMs expressing VEGF-C hampered pulmonary metastasis was ascribed to augmented VEGFR3 expression on tumor vessels and was associated with increased vessel normalization and functionality. These data were substantiated in a cohort of ER^+^ BC patients, showing that VEGF-C expressing TAMs correlated inversely with BC malignant grade and TMEM assembly. In summary, we propose a mechanism whereby TAM-derived VEGF-C attenuates pulmonary metastasis through mediating vascular normalization and is associated with increased VEGFR3 expression in tumor blood vessels and reprogramming of TAMs, both hampering cancer cell intravasation.

## Results

### Concurrent VEGF-C Expression in TAMs Correlates with Lymph Node Metastasis and Podoplanin Expression in TAMs

To elucidate whether TAMs control the route of metastatic dissemination of cancer cells, we employed two metastatic mammary cancer cell lines, 4T1 and 66cl4, which are derived from a single mouse mammary tumor cell type but exhibit different capabilities to spread to proximal and distant organs. ^17^ Both cell lines display features of human TNBC, *i.e.,* absence of estrogen and progesterone receptors, and amplification of *Erb-B2* (a gene homologous to Human Epidermal Growth Factor Receptor 2, *HER2*).

We ^18^ and others ^17^ have previously shown that 4T1 cancer cells preferentially disseminate to the lungs, while the 66cl4 cancer cells favor the lymph nodes. To confirm our previous findings, 4T1 and 66cl4 cancer cells from lungs and lymph node metastases 32 days after cancer cell injection were dissociated to a single cell suspension and cultured in the presence of the selection drug 6-thioguanine, for which both 4T1 and 66cl4 cancer cells display resistance. Clonogenic assays showed that 66cl4 cancer cells formed more colonies in the lymph node compared to lungs, while 4T1 cancer cells readily formed colonies from lungs and fewer colonies from the lymph nodes (**Figure 1A**), in line with previous reports of analogous experiments. ^18^ These findings suggest that 66cl4 cancer cells were more numerous in lymph nodes, whereas 4T1 cancer cells were preferentially located in lungs.

**Figure 1.**
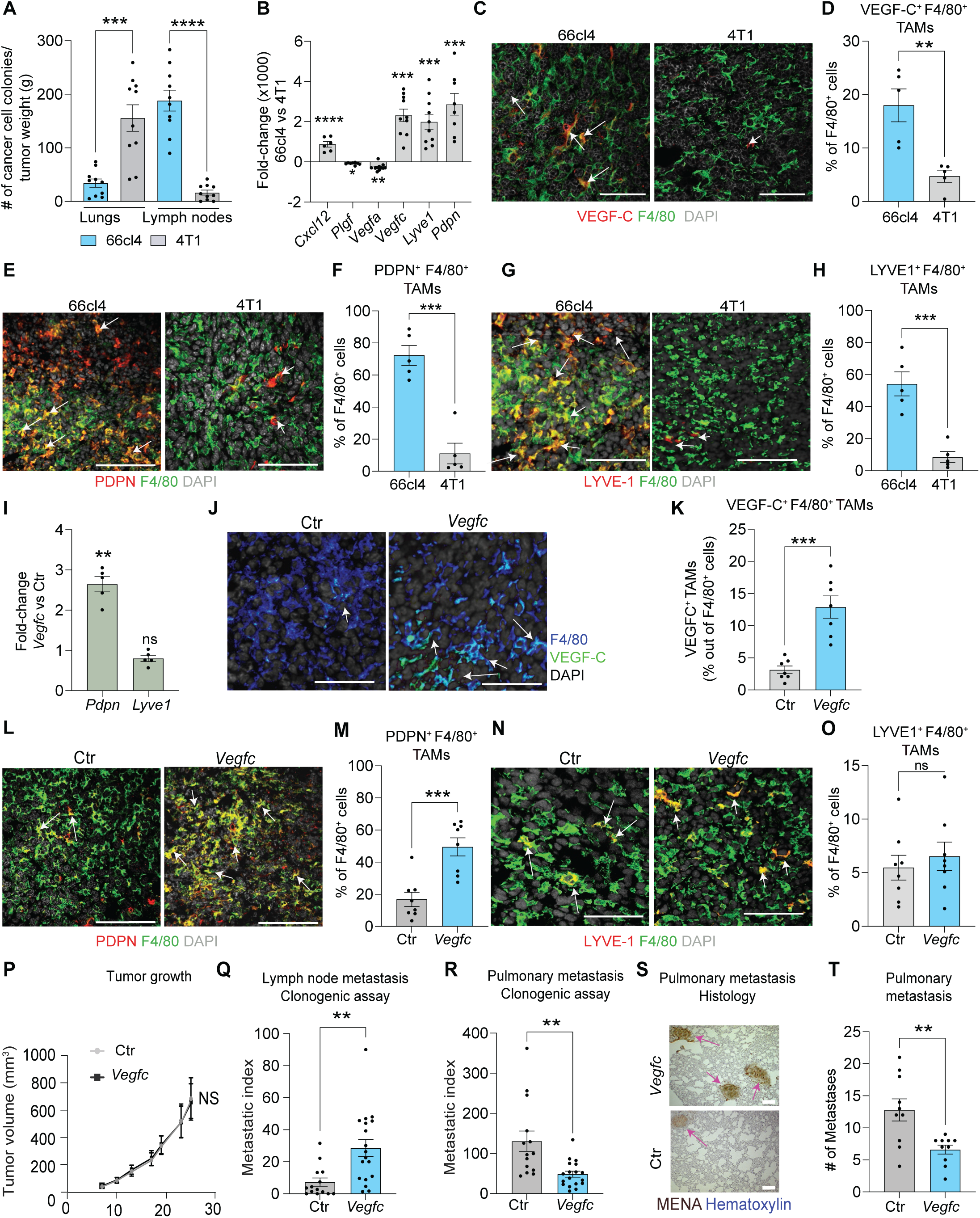
Concurrent VEGF-C Expression in TAMs Correlates with Lymph Node Metastasis and Podoplanin Expression in TAMs. (A-H) 66cl4 and 4T1 cancer cells were injected into the mammary fat pad of syngeneic mice. (A), Number of 66cl4 and 4T1 metastatic colonies in lungs and lymph nodes per gram of primary tumor weight (metastatic index). Statistical analyses were performed by Students t-test, stars indicate significance: ****p<0.0001; n=10 mice/group (B), Fold-change of the indicated genes in flow cytometry-sorted 66cl4-TAMs as compared to 4T1-TAMs, analyzed by qPCR. Statistical analyses were performed by Students t-test, stars indicate significance: *p<0.05, **p<0.01, ***p<0.001 and ****p<0.0001; n=6-10 mice/group (C–H), Immunofluorescence analysis of 66cl4 and 4T1 tumor sections immunoassayed for the pan-macrophage marker F4/80 (green) and cell nuclei (DAPI, grey). In (C,E,G), representative images for VEGF-C (red) in (C), PDPN (red) in (E), LYVE-1 (red) in (G). Morphometric analysis of VEGF-C (D), PDPN (F) and LYVE-1 (H) expressing F4/80^+^ TAMs are displayed in the graphs (D,F and H). Statistical analyses were performed by Data values are presented as mean ± SEM. Students t-test, stars indicate significance: **p<0.01, ***p<0.001; n=5 mice/group. At least 10 optical fields (OF) are visualized per tumor section and representative images are shown. Bars represent 50 μm. (I) BMDMs transduced with Ctr or *Vegfc*-LV and subjected to qPCR. Graph shows *Pdpn* and *Lyve1* gene expression. Students t-test, stars indicate significance: **p<0.01, ns indicated non-significant; n=5 (J–O) 4T1 cancer cells were injected into the mammary fat pad of syngeneic chimeric mice harboring haemopoietic cells transduced with *Vegfc* or Ctr LVs. Immunofluorescence analysis of 4T1-Ctr and 4T1-*Vegfc* tumor sections for F4/80 (blue), VEGF-C (green) and DAPI (grey) in (J), F4/80 (green), PDPN (red) and DAPI (grey) in (L) and F4/80 (green), LYVE-1 (red) and DAPI (grey) in (N). Morphometric analysis of VEGF-C expressing TAMs shown in (K), PDPN expressing TAMs in (M) and LYVE-1 expressing TAMs in (O). Students t-test, stars indicate significance: ***p<0.001; n=7-8 mice. At least 10 optical fields (OF) are visualized per tumor section and representative images are shown. Bars represent 50 μm. (P-T), 4T1 cancer cells were injected into the mammary fat pad of syngeneic chimeric mice transplanted with haemopoietic cells transduced with *Vegfc* or Ctr LVs. Graphs show tumor growth of Ctr and *Vegfc*-treated 4T1 tumors in (P), Lymph node (Q) and lung (R) colonies per primary tumor weight [(g) metastatic index] in (R). Immunohistochemical analysis of MENA expressing cancer cells in lungs from tumor bearing Ctr and *Vegfc*-chimeric mice are depicted in (S). Graph shows metastatic index [cancer cell foci/ tumor weight (g)] in (T). Students t-test, stars indicate significance: **p<0.01; n=10-14 mice/group. Bars represent 100 μm, white arrows indicate tumor cell foci.

To investigate the involvement of TAMs in regulating the cancer cell metastatic fate, F4/80^+^ TAMs were sorted from 4T1- and 66cl4-tumors using fluorescence-activated cell sorting (FACS) and subjected to qPCR for genes associated with lymphatic and blood vessel formation. 66cl4-infiltrating TAMs displayed several thousand-fold increases of the pro-lymphangiogenic molecules *Vegfc, podoplanin* (*Pdpn)* and the lymphatic vessel endothelial hyaluronic acid receptor 1 (*Lyve1)* and the angiostatic molecule *Cxcl12* as compared to 4T1-infiltrating TAMs (**Figure 1B**). Even though PDPN and LYVE-1 are mainly expressed on lymphatic vessels, it has recently been shown that PDPN-expressing TAMs drive lymphangiogenesis and lymph endothelial intravasation, ^19^ while LYVE-1 expressing TAMs maintain arterial homeostasis ^20^ and tumor angiogenesis. ^21^ Contrary to TAMs infiltrating 66cl4 tumors, TAMs infiltrating 4T1 tumors exhibited higher levels of the pro-angiogenic molecules VEGF-A and placental growth factor (PLGF) (**Figure 1B**). Further analysis using immunofluorescence stainings of tumor sections showed that the a high number of 66cl4-infiltrating TAMs expressed VEGF-C (**Figure 1C and 1D**), PDPN (**Figure 1E and F**), and Lyve-1 (**Figure 1G and 1H**) as compared to 4T1-infiltrating TAMs. We then investigated whether VEGF-C expression by TAMs led to upregulation of LYVE-1 and PDPN in an autocrine manner. To do this, we transduced bone marrow-derived macrophages (BMDMs) with lentiviral vectors (LVs) containing *Vegfc* under the transcriptional control of a CMV promoter (LVs without any DNA sequence downstream to the CMV promoter were used as control, *Ctr)*. We found that enforced expression of VEGF-C (~40-fold vs. *Ctr)* led to increased *Pdpn* levels but did not significantly affect *Lyve1* expression (**Figures 1I and S1A**). Suggesting that VEGF-C-sourced from TAMs may control the phenotype and function of other cell types, such as endothelial cells (ECs), rather than macrophages. Therefore, we next investigated if the abundance of VEGF-C^+^ TAMs was associated with increased lymphangiogenesis by analyzing the expression of CD31, a marker of blood vessels, and VEGFR3, a marker of lymph vessels, in 4T1- and 66cl4-tumor sections. Indeed, the 66cl4 tumors displayed more VEGFR3^+^CD31^low^ lymphatic vessels as compared to 4T1 tumors (**Figures S1B and S1C**). Unexpectedly, CD31^high^blood vessels in 66cl4 tumors expressed VEGFR3 to a higher extent than blood vessels in 4T1 tumors (**Figure S1D and S1E**). Even though VEGFR3 expression is mainly restricted to lymphatic vessels in the adult, its expression can be induced during pathological conditions such as tumorigeneses ^22^, and wound healing. ^23^ Furthermore, increased expression of VEGFR3 on blood vessels during embryogenesis has been suggested to control vascular permeability. ^24^ In summary, TAMs derived from a mammary tumor model, which preferentially disseminates via the lymphatic route and gives rise preferentially to lymph node metastases as compared to pulmonary metastases, expressed higher levels of VEGF-C, PDPN and LYVE-1 in contrast to TAMs isolated from a tumor model (4T1), which preferentially disseminates via the hematogenous route and gives rise preferentially to pulmonary metastases.

### 4T1 infiltrating TAMs expressing VEGF-C hampers pulmonary metastases

Building on these findings, we hypothesized that VEGF-C expression by TAMs could control the route through which cancer cells spread. Therefore, we generated chimeric mice harboring hematopoietic cells, including TAMs, overexpressing VEGF-C. To achieve this, we transduced hematopoietic stem/progenitor cells (HSPCs) with LVs, either empty (as indicated above) or driving the expression of *Vegfc*. Transduced HSPCs were then transplanted into recipient mice (hereon Ctr and *Vegfc*, respectively), which had previously undergone myeloablation through busulfan treatment. ^25^ After hematopoietic reconstitution (*i.e.,* after 6 weeks of HSPC transplantation), we injected 4T1 cells into the fat pad of the chimeric mice (**Figure S1F**).

We found an average of 1.5 integrated LV copies per mouse genome in bone marrow cells from both Ctr and *Vegfc*-chimeric mice, as determined by qPCR analysis (**Figure S1G**). By employing western blot analysis, we confirmed higher VEGF-C protein expression *ex vivo* in BMDMs isolated from transplanted *Vegfc* mice compared to control mice (**Figure S1H**). This observation coincided with a higher accumulation of 4T1-infiltrating TAMs expressing VEGF-C in the *Vegfc* group (**Figures 1J and 1K**). Importantly, the expression level of other molecules related to *Vegfc* such as *Vegfa, Vegfd*, VEGF family receptors (*Vegfr1*, *2*, and *3*) or the co-receptors Neuropilin (*Nrp1*) or *Nrp2* was not modulated by *Vegfc* overexpression (**Figure S1I**). In tumor sections, we found elevated levels of VEGF-C, which coincided with the accumulation of PDPN^+^ F4/80^+^ TAMs (**Figure 1L and 1M**) but not with LYVE1^+^ F4/80^+^ TAMs (**Figure 1N and 1O**). This observation is agreement with the results obtained upon enforcing VEGF-C expression in BMDMs (see **Figure 1I**).

Overexpression of VEGF-C preferentially in TAMs did not alter 4T1 tumor growth rate (**Figure 1P**) or the final tumor burden (**Figure S1J**) 30 days after cancer cell injection. Remarkably, clonogenic assays of lymph nodes and lungs from *Vegfc* chimeric mice implanted with 4T1 tumors, which preferentially give rise to lung metastases, showed increased cancer cell colonies in lymph nodes (**Figure 1Q**), but reduced colonies in the lungs (**Figure 1R**) or blood (**Figure S1K**) compared to controls. In agreement with this finding, immunohistochemical analysis of MENA-expressing cancer cells in the lungs from *Ctr* and *Vegfc*-chimeric mice implanted with 4T1 tumors confirmed that overexpression of *Vegfc* preferentially in TAMs decreased the incidence of pulmonary metastasis compared to the control (**Figures 1S and 1T**). In summary, our data indicate that the accumulation of VEGF-C^+^ PDPN^+^ TAMs can be involved in directing cancer cells to the lymph nodes rather than the lungs.

### TAM-derived VEGF-C normalizes the tumor vasculature

We then investigated whether *Vegfc* overexpression trough TAMs affected lymphatic and blood angiogenesis by analyzing tumor sections from *Ctr* and *Vegfc* chimeric mice stained for VEGFR3 and CD31. The enforced expression of VEGF-C through TAMs increased the number lymphatic vessels in border and inner areas of the tumor (**Figures 2A–2D**). Of note, VEGF-C enforced expression also increased the number of VEGFR3^+^ LYVE1^−^ CD31^high^ blood vessels compared to control (**Figure 2E**) while the number of CD31^high^ blood vessels objects were decreased (**Figure 2E**) as well as the CD31^+^ vessel area was decreased in *Vegf-c* 4T1 tumors as compared to controls (Figure S2A), which may suggest normalization of the vessels in the presence of VEGF-C expressing TAMs. In summary these results suggest that enforced expression of VEGF-C by TAMs may affect vessel functionality while promoting lymphangiogenesis.

**Figure 2.**
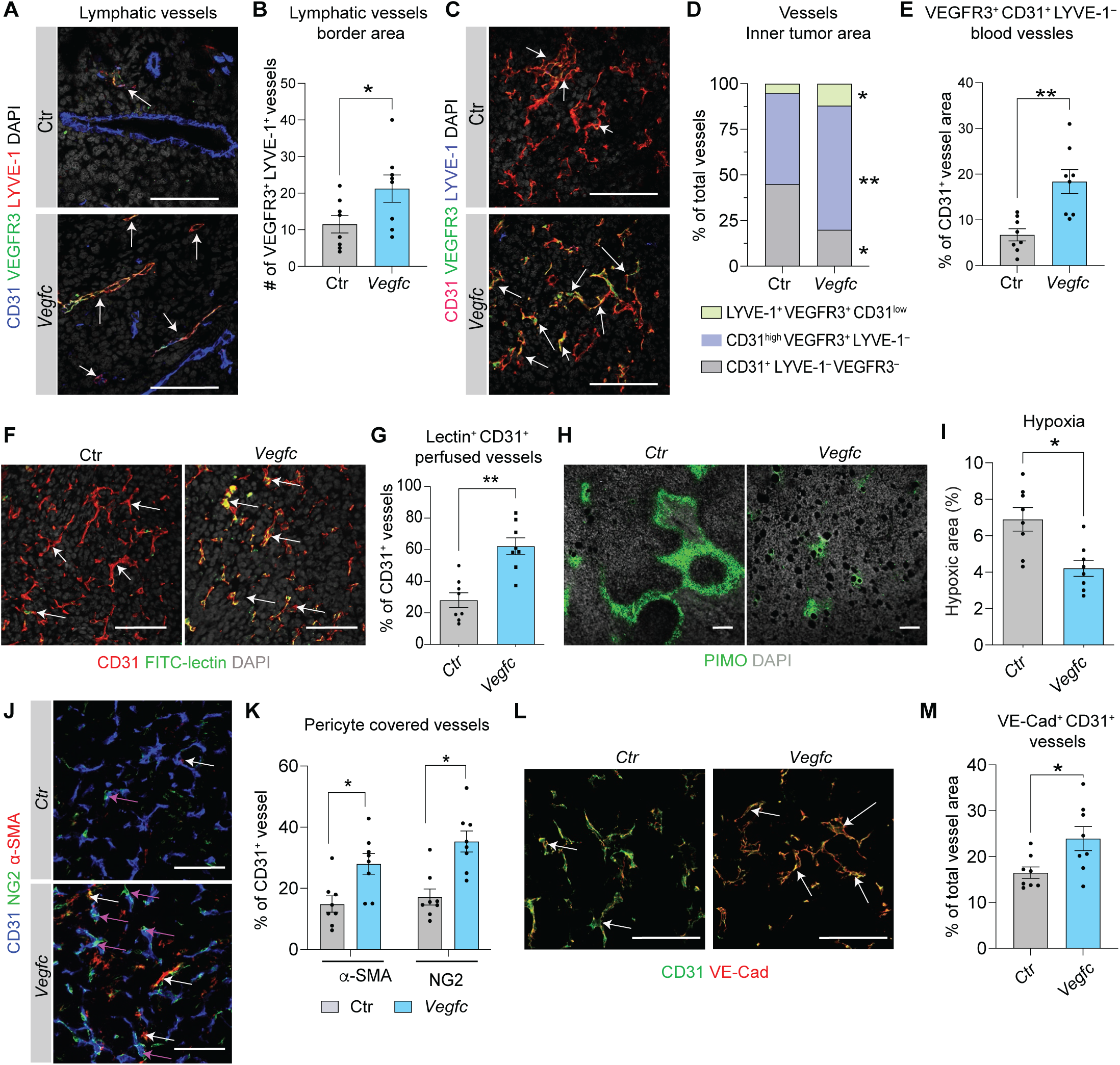
TAM-derived VEGF-C normalizes the tumor vasculature. (A-E) Immunofluorescence analysis of 4T1 tumor sections immunoassayed for CD31 (blue), LYVE-1 (red), VEGFR3 (green) and DAPI (grey) in (A). CD31 (red), LYVE-1 (blue), VEGFR3 (green) and DAPI (grey) in (B). Graphs shows number of VEGFR3^+^ LYVE-1^+^CD31^low^ lymphatic vessels in (B and D), number of VEGFR3^+^CD31^high^ vessel objects, CD31^high^ vessel objects, and CD31^low^ vessel objects (D), area of VEGFR3 expression in CD31^high^ blood vessels in E. Students t-test, stars indicate significance *p<0.05; **p<0.01, n=8 mice/group. All data values are shown as the mean ± SEM. Representative images are shown. At least 10 OFs are visualized per tumor section and representative images are shown. Bars represent 50 μm. White arrows indicate double positive cells. (F-I) 4T1 tumor bearing Ctr-and *Vegfc*-chimeric mice were injected with FITC-lectin intravenously and PIMO intraperitoneally before sacrificing. Representative images show immunostaining for CD31 (red), FITC-conjugated lectin (green) and DAPI (grey) in (F), PIMO (hypoxia) and DAPI (grey) in (H). Graph shows morphometric analysis of vessel perfusion in (G), hypoxia in (I). Students t-test, stars indicate significance *p<0.05; **p<0.01, n=8 mice/group. All data values are shown as the mean ± SEM. Representative images are shown. At least 10 OFs are visualized per tumor section and representative images are shown. Bars represent 50 μm. White arrows indicate double positive cells. (J-M) T Ctr- and *Vegfc*-tumor section were stained for CD31 (blue), the pericyte markers α-SMA (red) and NG2 (green), and DAPI (grey) in (J) and CD31 (green) and VE-Cadherin (red) in (L). Graph shows morphometric analysis of pericyte covered vessels in (K) and VE-Cadherin area out of CD31^+^ area in (M). Students t-test, stars indicate significance *p<0.05; n=8 mice/group. All data values are shown as the mean ± SEM. Representative images are shown. At least 10 OFs are visualized per tumor section and representative images are shown. Bars represent 50 μm.

Blood vessel normalization hampers cancer cell invasion and seeding of pulmonary metastases, ^10,15^ and is generally associated with improved vessel perfusion, increased pericyte coverage as well as decreased tumor hypoxia. ^10,26^ 4T1 tumor-bearing Ctr and *Vegfc* chimeric mice were injected with FITC-conjugated lectin, which will only label functional vessels with a blood flow, or the hypoxic marker pimonidazole hydrochloride (PIMO), intravenously or intraperitoneally respectively. Sections of 4T1 tumors from *Vegfc* chimeric mice displayed a significantly higher number of lectin perfused blood vessels (**Figures 2F and G**) and reduced tumor hypoxia (**Figures 2H and 2I**) compared to controls. In agreement with this, the number of blood vessels that were covered with α-SMA and NG2 positive pericytes was augmented in 4T1 tumor implanted in *Vegfc* mice compared to controls (**Figures 2J and 2K**). Further characterization of the tumor vasculature revealed that the junctional adhesion molecule VE-cadherin, which regulates vascular permeability and limits cancer cell intravasation ^27^, was more abundant in tumors from the *Vegfc* group than control (**Figures 2L and 2M**). Additionally, VEGFR2^+^ endothelial cells were more abundant in the *Vegfc* group than in controls, suggesting that VEGF-C released by TAMs promotes vessel normalization through activating the VEGFR2/VEGFR3 heterodimer (**Figures S2B and S2C**). These results suggest that VEGF-C expression by TAMs is associated with tumor blood vessel normalization.

### Targeted modulation of VEGF-C expression in TAMs rewires the metastatic destiny of 4T1 cells

To further investigate the impact of TAM-derived VEGF-C on tumor vessels, we performed an *in vivo* co-mingling assay involving cancer cells and BMDMs, as outlined in **Figure S3A**. In this assay, BMDMs were transduced either (i) with the *Vegfc* LV to enforce *Vegfc* expression, (ii) with a LV that drives the expression of a short hairpin RNA (shRNA) against *Vegfc* (sh-*Vegfc* LV) to downregulate endogenous *Vegfc*, (iii) with an empty LV (Ctr); or (iv) with a control shRNA (sh-Ctr) as controls (**Figure S3B**). Transduced BMDMs were then mixed with 4T1 cancer cells in matrigel plugs and then injected subcutaneously into syngeneic mice. The implants were harvested 8 days post-injection. Immunostaining revealed that *Vegfc*-expressing BMDMs increased the fraction of vessels labeled with FITC-conjugated lectin or VEGFR3 (**Figures 3A and 3B**). Conversely, sh*-Vegfc* BMDMs reduced the number of functional vessels, as indicated by lectin staining and decreased the fraction of vessels expressing VEGFR3 (**Figures 3C and 3D**). Additionally, *Vegfc* BMDMs reduced tumor hypoxia whereas sh*-Vegfc* BMDMs increased it compared to controls (**Figures 3E and F**).

**Figure 3.**
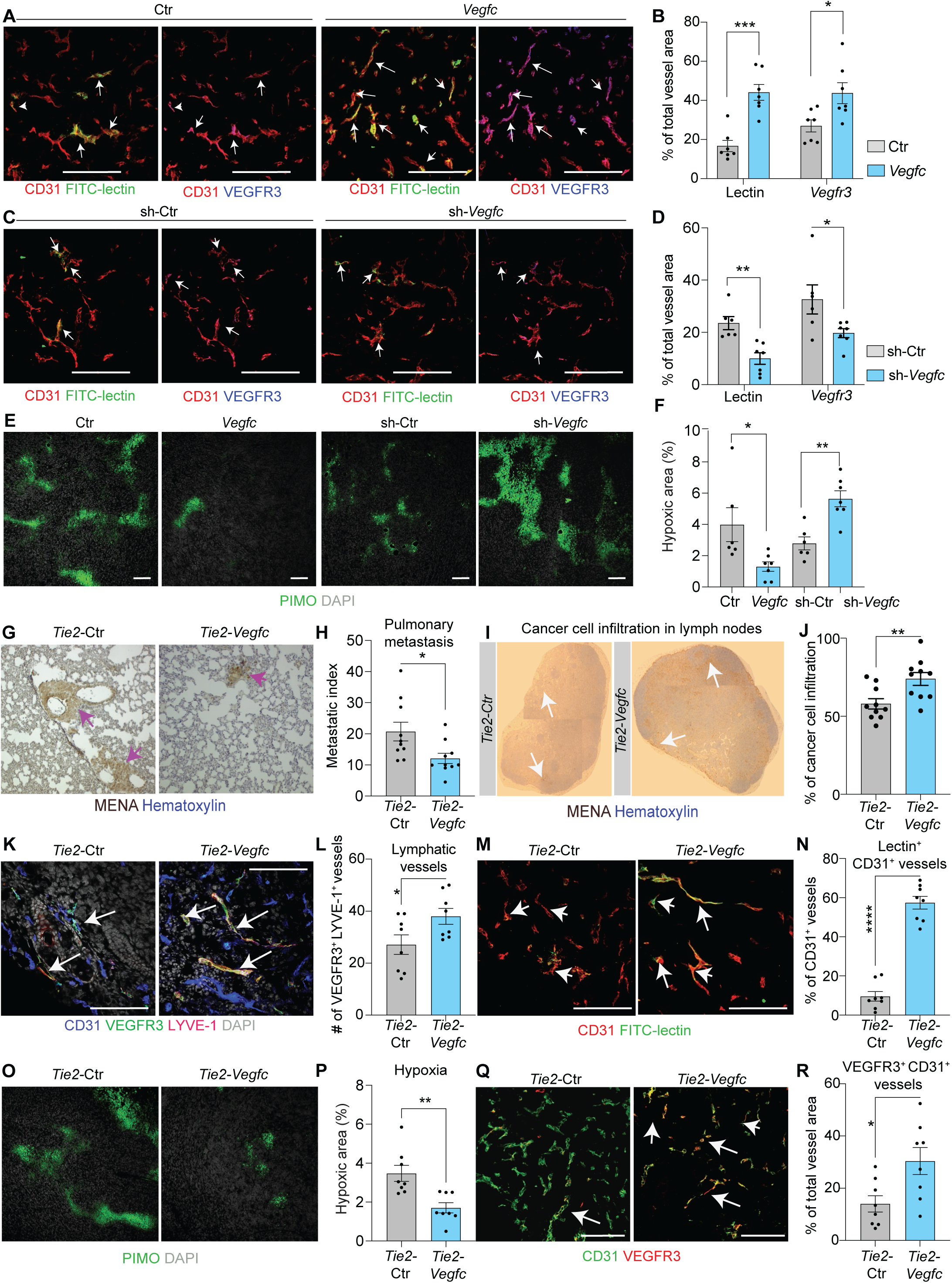
Targeted modulation of VEGF-C expression in TAMs rewires the metastatic fate of 4T1 cells. (A-F), 4T1 cancer cells were co-mingled in matrigel with BMDMs previously transduced with *Ctr*, *Vegfc,* sh-Ctr *or* sh*-Vegfc* LVs and injected subcutaneously into syngeneic mice. FITC-conjugated lectin and PIMO were injected before sacrification 8 days post-injection. Tumor sections were immunoassayed for CD31 (blue), FITC-conjugated lectin (green), VEGFR3 (red) and DAPI (grey) in (A and C), PIMO (green) and DAPI (grey) in (E). Morphometric analyses of perfused vessels that express VEGFR3 are depicted in (B and D), and hypoxic area in (F). Students t-test, stars indicate significance *p<0.05, **p<0.01, ***p<0.001; n=6-7 mice/group. All data values are shown as the mean ± SEM. Representative images are shown. At least 10 OFs are visualized per tumor section and representative images are shown. Bars represent 50 μm. 4T1 cancer cells were injected into the mammary fat pad of syngeneic chimeric mice harboring haemopoietic cells transduced with *Tie-*Ctr or *Tie2-Vegfc* lentiviral vectors in (G-R). (G-H) Immunohistochemical analysis of MENA expressing cancer cells in lungs from tumor bearing Ctr and *Vegfc*-chimeric mice are depicted in (G) and the graph shows metastatic index [cancer cell foci/ tumor weight (g)] in (H). Students t-test, stars indicate significance: *p<0.05; n=10. Bars represent 100 μm, white arrows indicate tumor cell foci. (I-J) Heamatoxylin and eosin staining of lymph nodes in (I). Cancer cells infiltration in lymph nodes are showed in (J) Students t-test, stars indicate significance: *p<0.05; n=10. Bars represent 100 μm. Cancer cell metastatic area is defined as the area of the lymph node containing MENA^+^ cancer cells. (K-L) Immunofluorescence analysis of *Tie2-Ctr and Tie2-Vegfc* tumor sections were immunoassayed for CD31 (blue), LYVE-1 (red), VEGFR3 (green) and DAPI (grey) in (K). Graphs shows number of VEGFR3^+^ LYVE-1^+^CD31^low^ lymphatic vessels in (L), p<0.05; n=7. Bars represent 100 μm, white arrows indicate tumor cell foci. (M-P) 4T1 tumor bearing *Tie2*-Ctr and *Tie2-Vegfc* chimeric mice were injected with FITC-lectin intravenously and PIMO intraperitoneally before sacrificing. Representative images show immunostaining for CD31 (red), FITC-conjugated lectin (green) and DAPI (grey) in (M), PIMO (hypoxia) and DAPI (grey) in (O). Graph shows morphometric analysis of vessel perfusion in (N), hypoxia in (Q). n=7 mice/group. All data values are shown as the mean ± SEM. Representative images are shown. At least 10 OFs are visualized per tumor section and representative images are shown. Bars represent 50 μm. (Q-R) Immunofluorescence analysis of *Tie2*-Ctr and *Tie2-Vegfc* tumor sections were immunoassayed for CD31 (blue), VEGFR3 (green) and DAPI (grey) in (Q). VEGFR3+ area of CD31^high^ blood vessels was assessed in (R). Students t-test, stars indicate significance *p<0.05, n=7 mice/group. All data values are shown as the mean ± SEM. Representative images are shown. At least 10 OFs are visualized per tumor section and representative images are shown. Bars represent 50 μm.

To further investigate the effects on tumor vasculature and metastatic seeding upon VEGF-C expression from TAMs, we decided to leverage on a LV incorporating the TIE2 promoter, which has been extensively used by us and others to convey molecular cargos selectively to tumors through TAM targeting ^28–30^. To this aim, we generated LVs containing either a *Vegfc* cDNA or non-coding DNA sequence (*Ctr*) downstream a *Tie2* promoter/enhancer sequence. HSPCs transduced with the resulting LVs were engrafted into busulfan-conditioned mice generating chimeric *Tie2-Vegfc* or Tie2-Ctr mice. Upon hematopoietic reconstitution 4T1 cells were injected into the mammary fat pad of these chimeric mice (**Figure S3C**). Mice were sacrificed 28 days after cancer cell injection. Although, we did not observe any effect on tumor weight (**Figure S3D**), *Tie2-Vegfc* reduced pulmonary metastasis, whereas promoted metastatic dissemination of cancer cells into lymph nodes, and concomitantly increased the number of lymph vessels in the primary tumors (**Figures 3K-L**). Furthermore, in agreement with our previous observations, *Tie2-Vegfc* increased vessel functionality (**Figure 3M and 3N**), which led to reduced hypoxia (**Figure 3O and 3P**) and vessel leakage (**Figures S3E and S3F**). Moreover, *Tie2-Vegfc* increased the expression of VEGFR3 in blood vessels (**Figure 3Q and 3R**). In summary, VEGF-C^−^expressed specifically by TAMs normalize blood vessels and increase vessel functionality, thus lowering the number of pulmonary metastases and redirecting cancer cells to the lymph nodes.

### VEGF-expressing TAMs do not alter the tumor immune composition

Lymphatic vessels are extensively involved in regulating immune cell trafficking, which may affect tumor growth and metastatic dissemination. ^4^ We therefore investigated if TAM-derived VEGF-C affected the tumor immune landscape by employing flow cytometry analysis. VEGF-C overexpression in TAMs did not affect the fraction of immune cells infiltrating tumors, including CD11b^+^ myeloid cells, CD11b^+^F4/80^+^ TAMs, Ly6G^+^ neutrophils, Ly6C^+^ monocytes, immunostimulatory, often referred as M1-like TAMs or immunosuppressive TAMs, often termed M2-like TAMs (**Figures S4A and S4B**). Immunostimulatory TAMs were identified by high expression of F4/80^+^, MHC class II, CD11c^+^, CD80^+^, CD86^+^ and MHC class I^+^ while immunosuppressive F4/80^+^ TAMs were identified by low expression of MHC class II or elevated levels of mannose receptor c-type 1 (MRC1). Moreover, TAM-driven VEGF-C did not affect the accumulation of T cells (neither total CD3^+^ T cells or CD4^+^ and CD8^+^ T cell subpopulations) or CD49b^+^ NK cells (**Figure S4C**). Furthermore, we found that *ex vivo* BMDMs engineered to express VEGF-C did not display altered expression levels of immunostimulatory cytokines such as C-X-C motif ligand (*Cxcl)-9, -10 and -11, interleukin 1a (Il-1a),* or immunosuppressive factors including *Il-10, arginase1* (*Arg1*), *Mrc1* and *transforming growth factor* (*Tgfβ)*, detected by qPCR (**Figure S4D**). Altogether, these observations indicate that TAM-derived VEGF-C does not alter the immune landscape of tumors as well as the immune activation or polarization status of TAMs. Hence, the differential metastatic behavior observed upon TAM-mediated VEGF-C expression may be attributed to a direct effect on tumor vessels.

### VEGF-C expressing TAMs inversely correlates to TMEM assembly and grade of malignancy in human breast cancer

Based on our experimental data, we speculated that perivascular TAMs expressing VEGF-C may promote vessel normalization in human tumors and, therefore, be more frequent in low-grade BC as compared to high-grade BC. The majority of available TNBC patient specimens belong to high-grade tumors (*i.e.,* grade 3, where the cancer cells are poorly differentiated and undergo rapid proliferation), whereas for our study we needed a population of specimens comprising both low grade and high-grade tumors. Building on this, we focused on estrogen receptor (ER)-positive breast cancer, which is also the most prevalent type in BC patients (~70 %) and can be targeted with endocrine therapy. However, half of the treated patients will develop metastatic disease within 20-years of diagnosis ^31^. Hence, we evaluated the accumulation of bulk and perivascular TAMs by immunostaining tumor sections from ER^+^ BC patient specimens of different histological grades (Nottingham Histological Grade I-III, where grade I and II indicate low grade, and grade III accounts for high grade). Both the total amount of CD68^+^ TAMs (**Figures 4A and 4B**) and perivascular TAMs (**Figure 4C**) accumulated to a higher extent in high-grade ER^+^ BC as compared to low-grade ER^+^ BC. Additional immunofluorescence staining, which also included VEGF-C, revealed that both bulk TAMs (**Figures 4D and 4E**) and perivascular TAMs (**Figures 4D and 4F**) expressed VEGF-C to higher levels in low-grade than in high-grade ER^+^ BC. Whereas no differences were observed in the number of blood vessels (**Figure S4E**).

**Figure 4.**
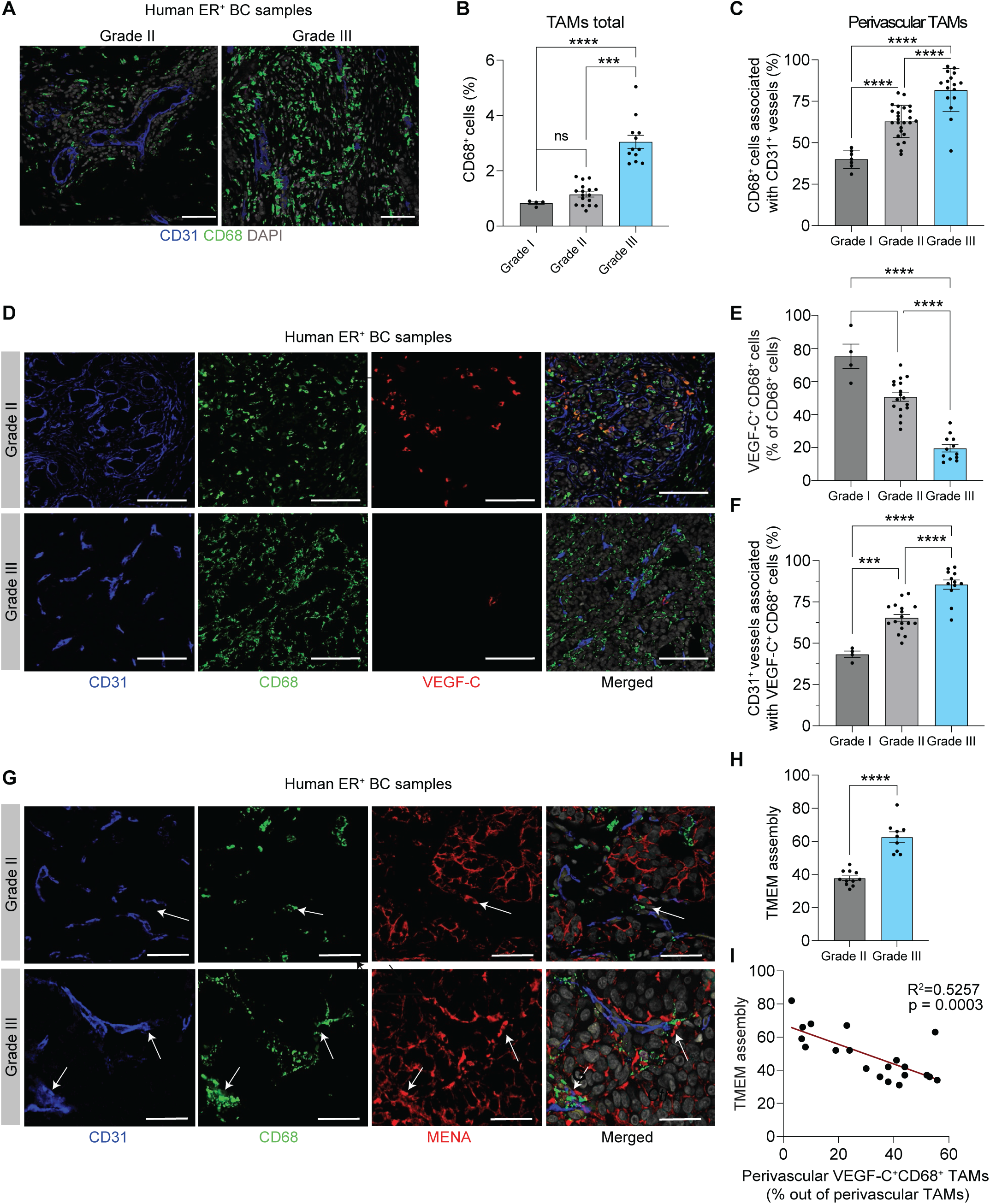
TAM-derived VEGF-C inversely correlates to the grade of malignancy in human breast cancer and density of TMEM. (A–C), Immunofluorescence analysis by confocal microscopy of human ER^+^ breast tumors immunoassayed for CD31 (blue); pan-macrophage marker CD68 (green) and nuclei (DAPI; grey). Representative images in (A), quantitative analysis of CD68^+^TAMs in (B) and CD68^+^ perivascular TAMs in (C). (D–F), Immunofluorescence analysis by confocal microscopy of human ER^+^ breast tumors immunoassayed for CD31 (blue), CD68 (green), and VEGF-C (red). Representative images in (D) and quantitative analysis of VEGF-C expressing bulk TAMs in (E) and perivascular TAMs in (F). (G-I), Representative pictograms display TMEM assembly (white arrows). Whole BC specimens were immunoassayed for CD31 (blue), CD68 (green), MENA (red) and DAPI (grey), in (G). TMEM assembly was assessed by quantifying 20 high-magnification fields/tumor section, visualized in (H), Pearson’s correlation is (R^2^=-0.7250, **p<0.01) for TMEM assembly versus VEGF-C-expressing TAMs depicted in (I). Statistical analyses were performed by one-way ANOVA in (B,C,E and F) and by Mann-Whitney U test in (H); **p<0.01 ***p<0.001, ****p<0.0001; n=4 for grade I, n=14 for grade II and 10 for grade III. Data are presented as the mean ± SD. At least 20 OFs are visualized per tumor section and representative images are shown. Bars represent 100 μm in (A,D) and 20 μm in (G).

As information for distant metastases for this patient cohort was not available, we evaluated the occurrence of TMEM assemblies, which is a clinically validated prognostic marker for the occurrence of distant metastasis in BC patients. ^12,32,33^ Immunofluorescence staining of TMEM assemblies revealed that the TMEM score was increased in high-grade compared to low-grade ER^+^ BC tumors (**Figures 4G and 4H**). Importantly, the TMEM score was inversely correlated with the presence of TAMs expressing VEGF-C (**Figure 4I**), and thus TAMs expressing VEGF-C may also inversely correlate with lung metastases. Altogether, these results suggest that VEGF-C expression by TAMs may play a key role in determining the preferential route of cancer cell seeding for distant metastases. Indeed, VEGF-C expression by TAMs may normalize tumor vessels limiting cancer cell spreading through the hematogenous route favoring the lymphatic route. Thus, our work points out a new molecular mechanism by which TAMs may foster or limit lung metastases in BC and sets the ground for identifying new prognostic tools and designing targeted interventions aimed at interfering with the metastatic process.

## Discussion

TAMs are regulators of tumor progression and have been linked to poor clinical outcomes in most cancers. ^34^ Like macrophages in many other tissues, TAMs show remarkable functional plasticity and often exhibit a spectrum of activation states that span from an immunostimulatory and anti-metastatic phenotype (sometimes referred to as M1-like) to an immunosuppressive and pro-metastatic phenotype (also referred to as M2-like). ^35^ TAMs typically express markers associated to both polarization states and their functional roles are, in part, determined by their spatial localization ^36^. For instance, perivascular macrophages promote angiogenesis by secreting factors such as VEGF-A, which increases vessel permeability and favors cancer cell intravasation. ^11,13^ Perivascular TAMs are furthermore a part of the TMEM assembly that functions as gateways for hematogenous spreading. ^11–14,32,37^

For the past decades it has been debated to what extend metastatic cancer cells in lymph nodes give rise to metastases in other distant organs, such as lungs, brain or liver; or whether distant metastases originate directly from the primary tumors ^4^. As VEGF-C is the main factor stimulating lymphangiogenesis, it has been correlated to lymph node metastasis. ^3^ VEGF-C stimulates lymphangiogenesis via activation of VEGFR3 but can also regulate angiogenesis by binding to VEGFR2/VEGFR3 heterodimers (expressed by blood vessels during embryogenesis) or endothelial tip cells in tumor blood vessels. ^38,39^ It has been reported that VEGFR3 signaling can tune the proportion of tip vs stalk endothelial cells, preventing excessive angiogenic sprouting. ^39^ In contrast, deletion of VEGFR3 in blood vessels enhances VEGFR2 expression that stimulates the VEGFA/VEGFR2 pathway, resulting in excessive angiogenesis in Lewis Lung Carcinoma. ^39^ Previous studies have demonstrated that supraphysiological overexpression of VEGF-C in the primary tumor leads or in transgenic mice increases the occurrence of metastases in both lymph nodes and lungs ^40,41^. In agreement with observations, by using photoconversion of mammary cancer cells, it has been showed that cancer cells from lymph node metastases can colonize the lungs. ^6^ However, further studies are necessary to determine the precise fraction of cancer cells in lung metastases originating from lymph nodes or primary tumor. More recent studies leveraging on clonal evolution of lung metastases in cancer patients suggest that most distant metastases in the majority of patients arise from the primary tumor. ^8,42^. We therefore hypothesize that physiological levels of VEGF-C, secreted by macrophages, promote vascular normalization and lymph angiogenesis, which prevents cancer cell intravasation and promotes lymph node metastases.

In line with this observation, our data do not only suggest that VEGF-C expressing TAMs are key regulators of functional vascular development but also redirects cancer cells to disseminate via lymphatic vessels instead of the hematogenous route by also inducing expression of PDPN. In line, Bieniasz-Krzywiec and colleagues suggested that PDPN expression by TAMs facilitated extracellular matrix degradation promoting lymphoinvasion facilitating lymph node metastases. ^19^ Here, we suggest that perivascular TAMs, expressing a combination of VEGF-C and PDPN, on one hand normalized the tumor vasculature and on the other hand facilitated lymphoinvasion by breast cancer cells. Studies on other tumor mouse models may be necessary to investigate the broader applicability of our observations. Further clinical relevance for VEGF-C-expressing TAMs in counteracting hematogenous metastatic dissemination was showed in a cohort of ER^+^ BC specimens. Perivascular VEGF-C^+^ TAMs correlated inversely with malignant grade and with the occurrence of TMEM complexes. Thus, even though VEGF-C levels in primary tumors and in serum correlate to malign tumors and lymph node metastases, ^3,4^ these levels might just be an indicative of malign disease rather than VEGF-C itself fueling the occurrence of distant metastases. Collectively, our data show that TAM-derived VEGF-C plays an unexpected role in normalizing the tumor vasculature, redirecting cancer cells to preferentially spread to the lymph nodes and dampening pulmonary metastases. This study paves the way for future works aimed at dissecting the contribution of VEGF-C in seeding of distant metastases from other tumor types, as well as identifying new prognostic markers and designing new targeted interventions aimed at thwarting the metastatic process.

## Materials and Methods

### STAR★Methods

#### Key resources table-

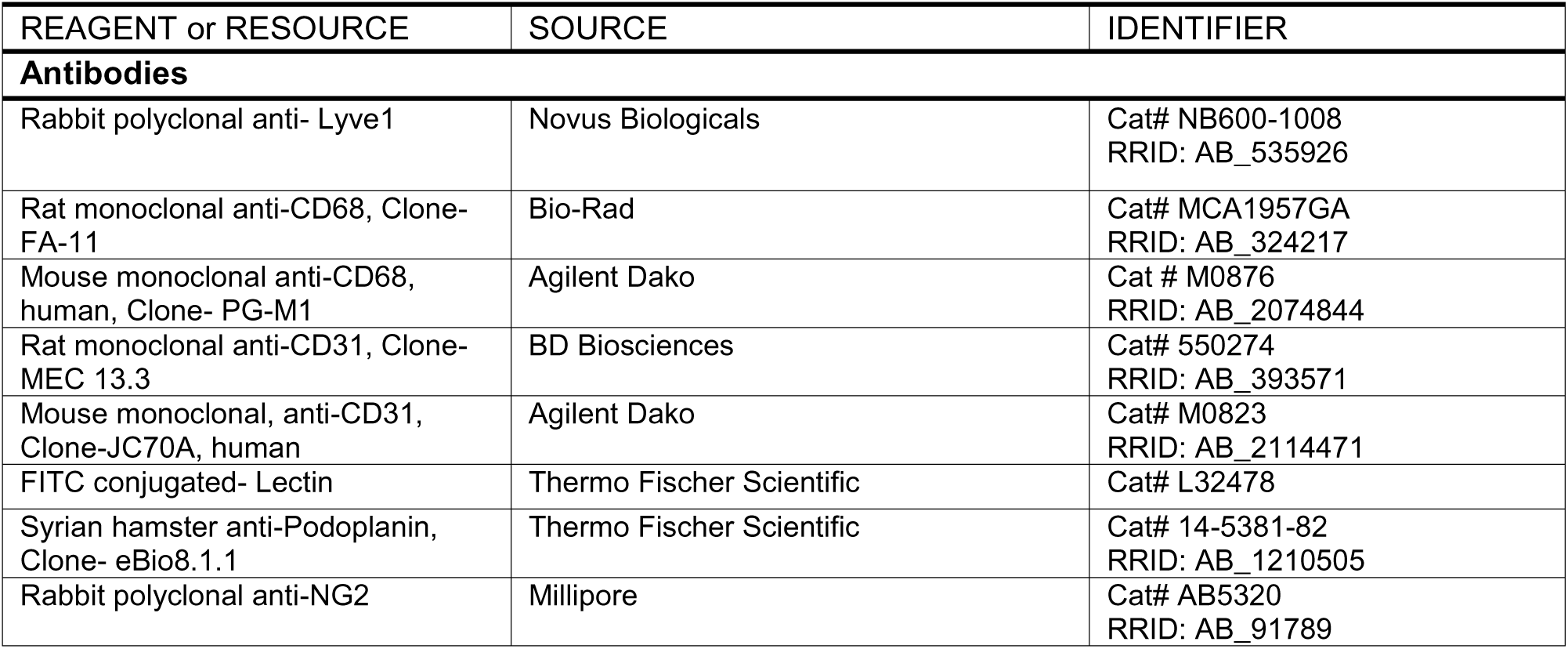
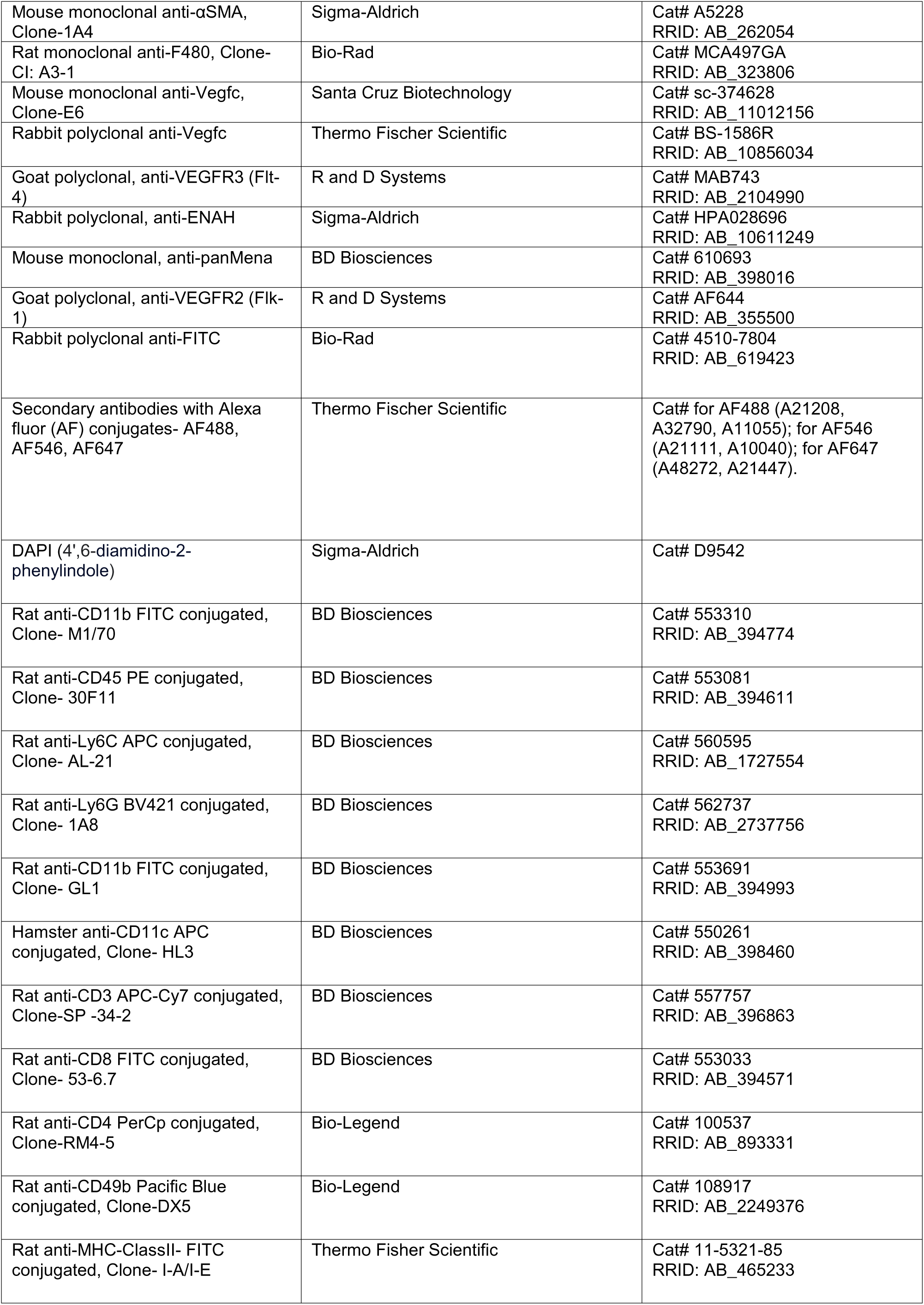
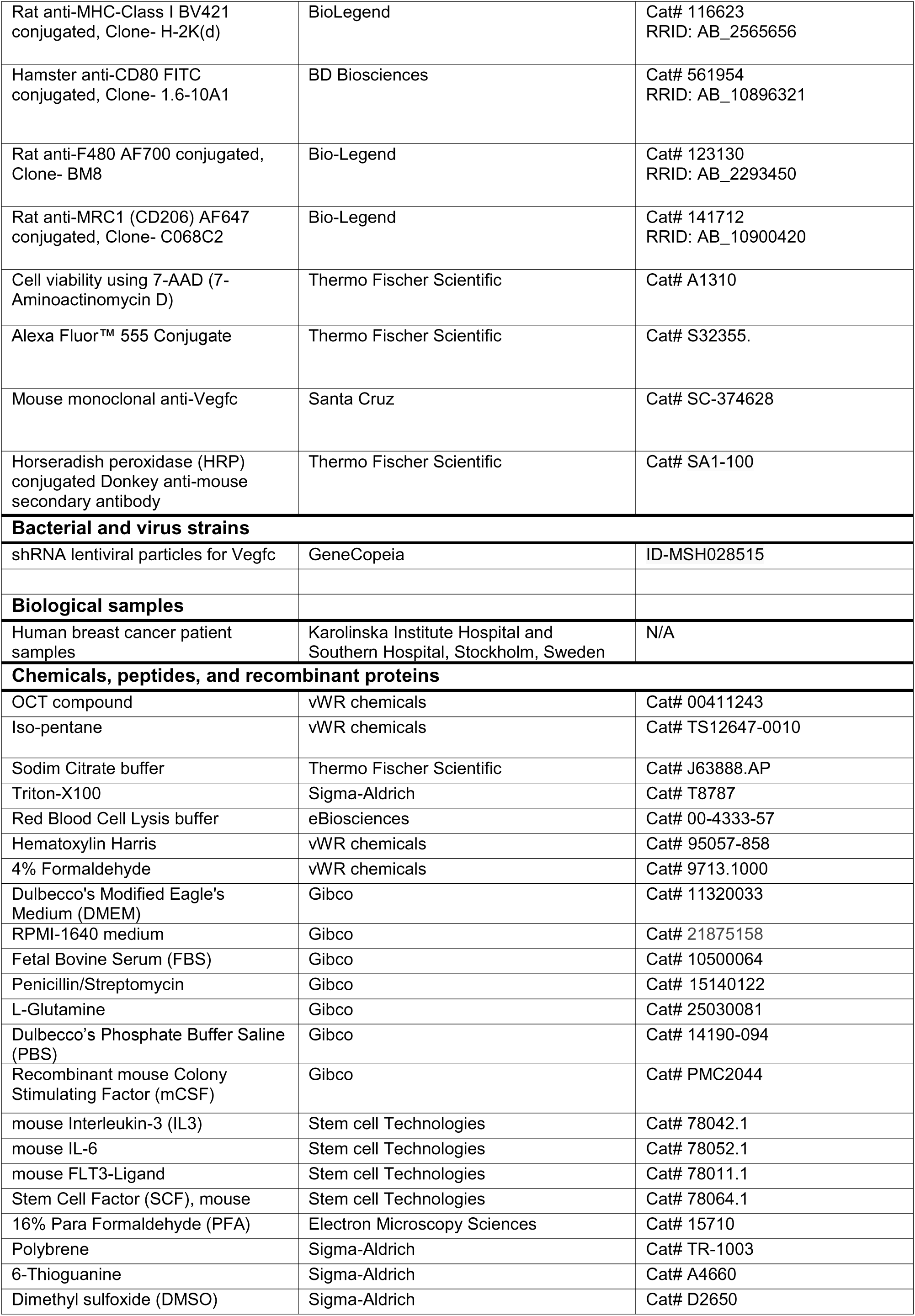
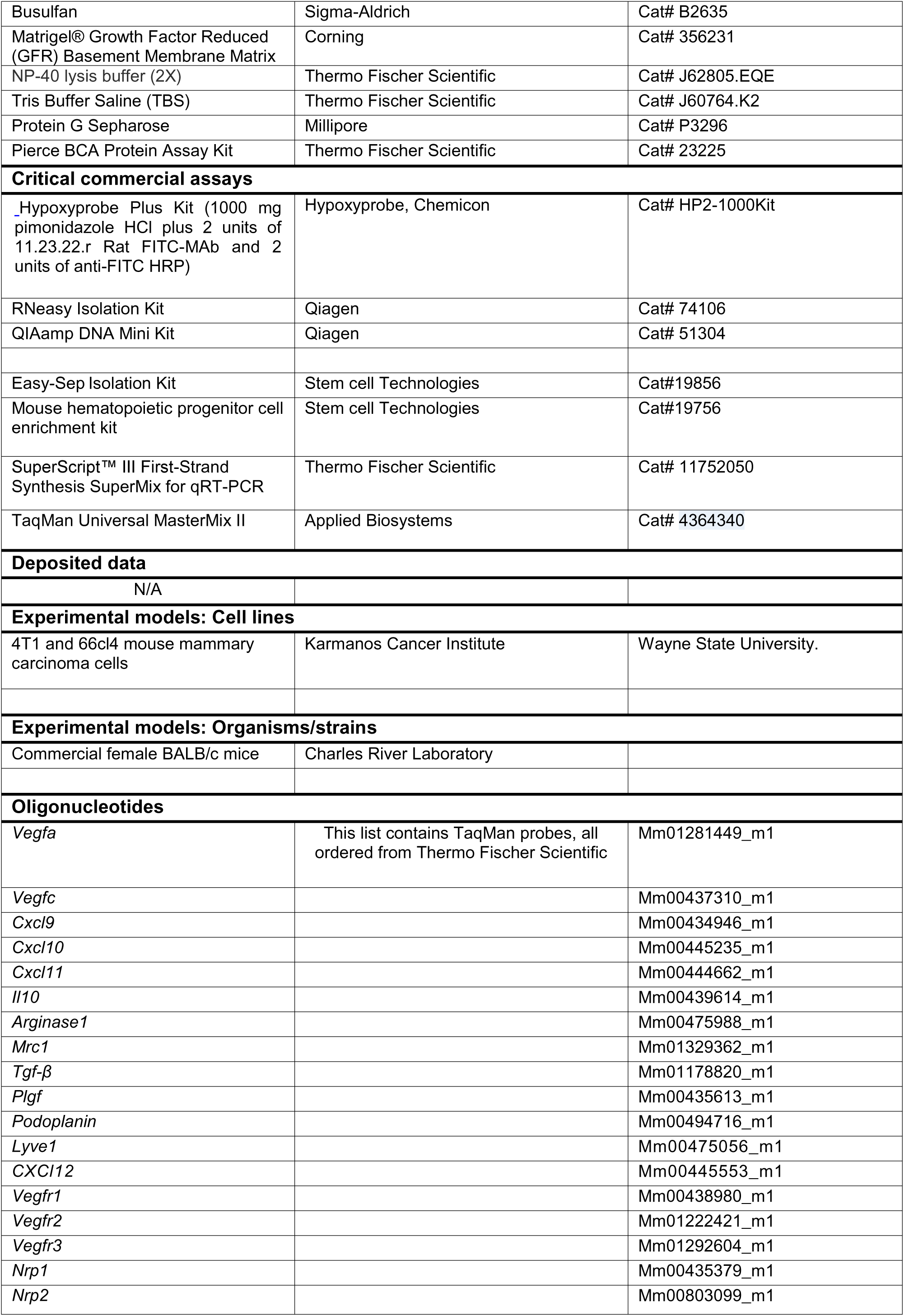
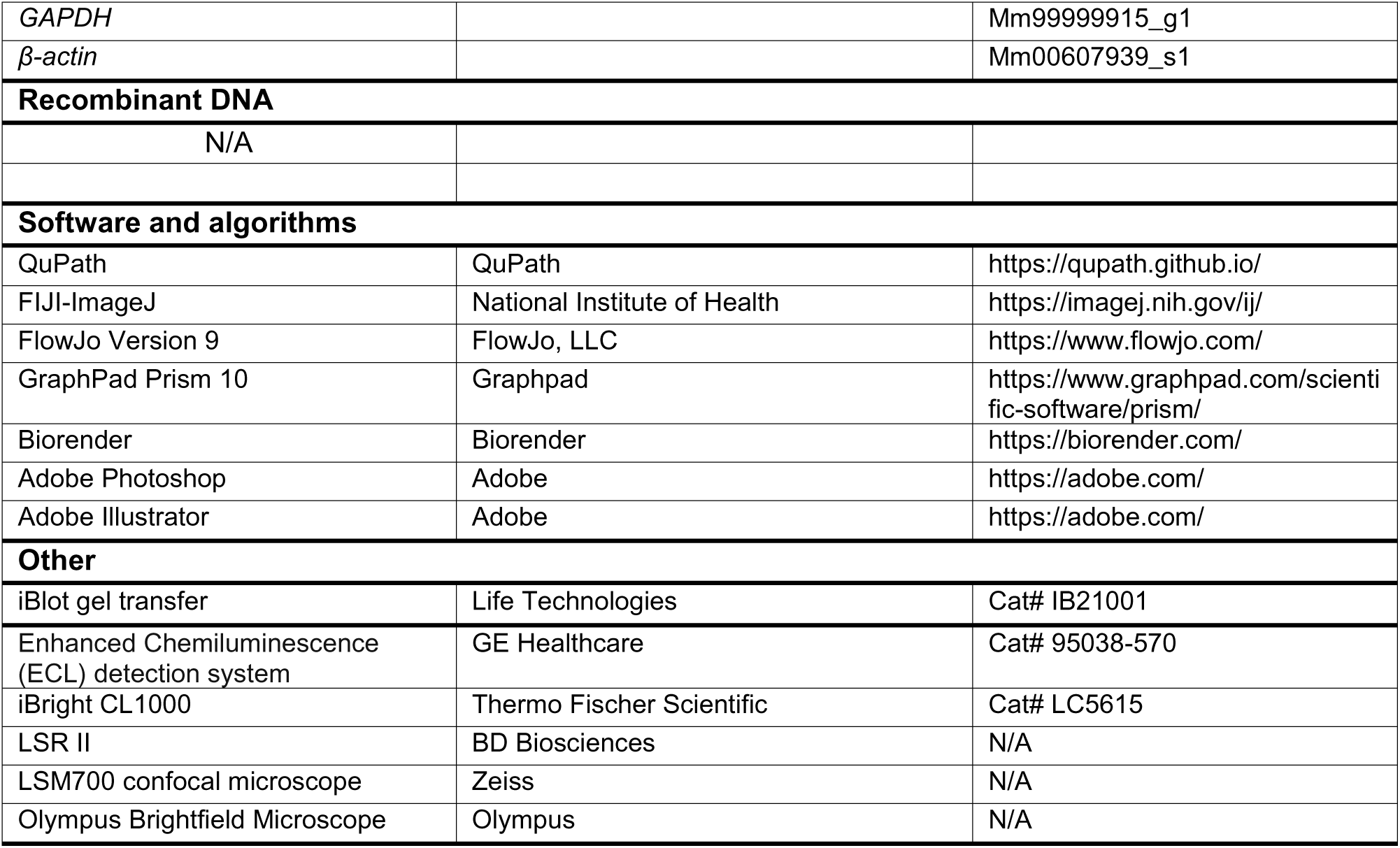

#### Lead contact

Further information and requests for resources and reagents should be directed and fulfilled by the corresponding contact, Charlotte Rolny and Mario Squadrito.

#### Cell culture

##### Tumor cell lines

HEK293T cells were grown in Dulbecco’s Modified Eagle’s Medium (DMEM) while 66cl4 and 4T1 mammary carcinoma cells were grown in RPMI-1640 medium (Gibco). Both the RPMI-1649 and the DMEM mediums were supplemented with 10% fetal bovine serum, 1% penicillin/streptomycin (Gibco) and 1% L-glutamine (Gibco). 4T1 and 66cl4 mouse mammary carcinoma cells were originally derived from a single spontaneous tumor that arose in a BALB/cfC3H mouse and were purchased from the Karmanos Cancer Institute at Wayne State University.

##### Generation of 4T1 cells expressing VEGF-C

1 x 10^6^ 4T1 cells were transduced with 1 x 10^8^ TU/ml of empty vector or *Vegfc* LVs in the presence of polybrene (Sigma-Aldrich).

##### Bone-marrow derived macrophages (BMDMs)

Bone marrow cells were acquired by flushing the femurs and tibias of female BALB/c mice as described before. ^43^. Cells were cultured in RPMI-1640 medium (Gibco) supplemented with 10% fetal bovine serum, 1% penicillin/streptomycin (Gibco), 1% L-glutamine (Gibco) and 50 ng/ml M-CSF (Gibco) for 8 days as described before. ^43^

#### Tumor experiments in mice

Female BALB/c mice (4-6 weeks old) were purchased from Charles River Laboratory. 2 x 10^5^ 4T1 cells were injected into the 4^th^ right mammary fat pad of anesthetized animals. Tumor size was measured externally using calipers, and tumor volumes were estimated using the following equation: V_sphere_ = 4/3 π x (d/2) 2 x D/2 (d: minor tumor axis, D: major tumor axis). Mice were euthanized at maximum 1000 mm^3^, and tumors were weighed after dissection. All ethical permits were obtained from the Swedish Board of Agriculture (N65/16 and 8525-2020).

#### Patient material

Paraffin-embedded whole tumor sections from breast cancer patients were obtained from Karolinska University Hospital and Southern Hospital, Stockholm, Sweden. All breast cancer patients at the Karolinska University Hospital and Southern Hospital have approved and signed a written consent for biobanking, including research purposes that have been approved by the local ethical committee (2016/957-31; *ex vivo* tumor models).

### Method details

#### Generation of the VEGF-C vector

*Vegfc* ORF clone was purchased from Thermo Scientific Fisher and amplified by PCR (95 °C for 5 min, 5 cycles at 95 °C for 30 sec, 45 °C for 30 sec, 72 °C for 30 sec and 30 cycles at 95 °C for 30 sec, 68 °C for 30 sec, 72 °C for 30 sec).

Primers for *Vegfc* amplification are FW 5’TAAAGACCGGTCAAAAGTTGCGAGCC-3’ and RW as 5’-TCGGGGTCGACTACAATTTTCATTTTATTTTAAAC-3’ primers. The DNA coding for *Vegfc* was then inserted into a CMV.GFP.WPRE lentiviral vector sequence as described previously ^44^ downstream to the CMV promoter.

#### Generation of the *Tie2*-*Vegfc* LC

For the generation of the of the *Tie2*-*Vegfc* LV, we started from the previously described *hTie2*.GFP lentiviral vector sequence ^29^ containing *Tie2* promoter/enhancer sequences, and inserted 4 tandem copies of the sequence 5’-CGCATTATTACTCACGGTACGA-3’, a sequence complementary to the miR-126 ^29^ followed by a WPRE sequence by using the restriction enzyme sites SalI and ScaI. Next, we inserted a DNA sequence coding for VEGF-C downstream to the *hTie2* promotor replacing the GFP sequence by using the restriction enzyme sites BamHI and SalI generating the *Tie2*-*Vegfc* LV.

#### Generation of the sh-*Vegfc* and sh-Ctr LVs

To originate the sh-*Vegfc* LV we inserted the sequence 5’ACCGCCCAAGTCTGTGTTTATTGAAACGCGTTTCAATAAACACAGACTTGGGTTTTTTCGA-3’, a sequence coding for a shRNA targeting VEGF-C as previously described ^45^ expressed under a human U6 promotor, in a LV backbone. As reporter, we used an eGFP encoding sequence expressed under a human *PGK* promotor and followed by a WPRE sequence. Sh-LV vector was generated using a non-targeting control DNA sequence (SHC016 Sigma Aldrich; shCTR) as previously described ^43^.

#### Lentivirus production

Vesicular stomatitis virus-G protein (VSV-G)-pseudotyped lentivirus vectors either empty or encoding *Vegfc or sh-Vegfc,* were produced by transient four-plasmid co-transfection into HEK293T cells and concentrated using ultracentrifugation as described previously. ^46^ Lentiviral vector titers were determined on HEK293T cells by TaqMan analysis as described previously ^46^. Lentiviral vectors were resuspended in PBS and stored at −80 °C.

#### gDNA isolation and quantitative RT-PCR

To evaluate the integration of the viral backbone into the hematopoietic progenitor cells, bone marrow of transplanted tumor bearing CTR and VEGF-C mice was isolated and gDNA was purified using QIAmp DNA Mini Kit (Qiagen) as described. ^47^. Vector integration was determined by quantifying copy numbers of HIV normalized to housekeeping gene beta actin by quantitative RT-PCR (95 °C for 10min, 40 cycles at 95 °C for 15 sec and 60 °C for 1 min).

#### Quantitative PCR

Flow sorted 66cl4- and 4T1-tumor cells, 66cl4 and 4T1 infiltrating TAMs and BMDMs were collected in RLT buffer (Qiagen) and homogenized using a syringe with a 20G needle. RNA was extracted using the RNeasy Mini Kit (Qiagen) according to the manufacturer’s instructions and reverse-transcribed with the SuperScript™ III First-Strand Synthesis SuperMix for qRT-PCR kit according to the manufacturer’s instructions (Thermo Fischer Scientific). Genomic DNA was eliminated using gDNA Wipeout Buffer (Qiagen) for 2 min at 42 °C. qPCR was performed using either TaqMan Universal MasterMix II and TaqMan Gene Expression Assays (Applied Biosystems) in a total volume of 20μL. The polymerase was activated at 95 °C for 10 min and the PCR was performed for 40 cycles (95 °C for 15 s and 60 °C for 60 s). The list of genes and its corresponding TaqMan probe details are mentioned above.

#### Bone marrow transplant

Bone marrow cells were acquired by flushing the femurs and tibias of 6- to 8-week-old female BALB/c mice and treated with red blood cell lysis buffer (Sigma). Lineage negative hematopoietic stem and progenitor cells (HSPCs) were purified using EasySep Isolation Kit (StemCell Technologies) according to manufacturer’s instructions. 2 x 10^6^ lineage negative cells/ml were transduced with 1 x 10^8^ TU/ml of empty vector or *Vegfc* lentivirus in serum-free Stem-Span medium (StemCell Technologies) supplemented with 10 ng/ml IL-3, 20 ng/ml IL-6, 100 ng/ml SCF and 10 ng/ml Flt3-L (Stem Cell Technologies). 1 x 10^6^ transduced HSPCs were resuspended in PBS and injected into the tail vein of 6-week-old female Balb/c mice, that had undergone myeloablation using 20 mg/kg Busulfan (Sigma) by intraperitoneal (i.p.) injection on three consecutive days. Busulfan was solubilized in DMSO and diluted to 5 mg/ml in PBS. On the fourth day 12–14 × 10^6^ BM cells were given by tail vein injection. This procedure resulted in ~80 % chimerism. ^25^

#### Metastasis assay

##### Clonogenic assay

Lungs and inguinal draining lymph nodes were dissected into an enzymatic buffer, dissociated to single cell suspensions and cultured in RPMI-1640 (complete medium) supplemented with 60 μM 6-Thioguanine (Sigma-Aldrich) as previously described (18) ^18^. Colonies formed by tumor cells resistant to 6-Thioguanine were fixed with 4 % Formaldehyde (VWR Chemicals) and stained with Hematoxylin Harris (VWR Chemicals). Metastasis colonies were counted, and the metastatic index was calculated by using the equitation M_index_ = n/w (n: number of metastases, w: weight of tumor). ^18^

##### Immunohistochemistry

All tissue were extracted and immersed in 10% formalin in a volume ratio of tumor to formalin of 1:7. Tissues were fixed for 24 hours and embedded in paraffin, then processed for histological examination. 5 µm lung tissue was stained for hematoxylin and anti-pan Mena antibody (610693; BD Biosciences) wile lymph nodes were only subjected to hematoxylin and eosin staining. Mena+ metastasis foci were counted for the lungs while cancer cell infiltration into the T cell margin of the out of total lymph node was assessed by ImageJ.

#### Flow Cytometry

Single cell suspensions tumors were pre-incubated with anti-CD16/32 mAb (BioLegend) for 15 min on ice to prevent non-specific binding, with subsequent incubation with specific antibodies for 30 min on ice. Cells were stained using the following antibodies: CD11b (M1/70), CD45 (30-F11), Ly6C (AL-21), Ly6G (1A8), CD86 (GL1), CD11c (HL3), CD3 (500A2/145-2c11), CD8 (53-6.7), CD4 (RM4-5), CD49b (HM ALPHA2), MHC class II (I-A/I-E), MHC class I (H-2K(d)) and CD80 (16-10A1; all from BD Bioscience), F4/80 (BM8) and MRC1 (C068C2; BioLegend). The viability of cells was verified using 7-AAD (Life Technologies). Samples were acquired with a LSR II (BD Biosciences) and analyzed using Flow-Jo software (Tree Star).

#### Co-mingling assay

BMDM were transduced with 1 x 10^8^ TU/ml of empty vector, *Vegfc*, *shCtr* or *shVegfc* lentivirus vectors, 4 days post-differentiation. These transduced BMDMs were then embedded in 200 μL of growth factor reduced Matrigel (BD Bioscience) together with 2 × 10^5^ 4T1 tumor cells, 2 days after transduction, and injected subcutaneously into the flank of 8-week-old female BALB/c mice for 8 days.

#### Western Blotting

Cells were lysed in NP-40 (Thermo Fisher Scientific) with freshly added Halt™ Protease Inhibitor Cocktail (Thermo Fisher Scientific). Protein concentrations were determined using the bicinchoninic acid (BCA) method (Thermo Fisher Scientific) according to the manufacturer’s instructions. 20 μg of protein was incubated with antibodies against VEGF-C (Santa Cruz) for 2 h on ice before mixing with protein G-Sepharose (Millipore). The antibody complex was washed three times in lysis buffer and once in Tris-buffered saline (TBS). The beads were resuspended in sample buffer, boiled for 5 min, and subjected to electrophoresis in 10% or 4–12% Bis-Tris gels and transferred to nitrocellulose membranes using iBlot2 (Life technologies). Membranes were probed for VEGF-C followed by horseradish peroxidase-conjugated secondary antibodies (Thermo Fischer Scientific). Membranes were then washed and visualized with an enhanced chemiluminescence (ECL) detection system (GE Healthcare) and iBright CL1000 (Thermo Fisher Scientific). Signals were quantified using ImageJ software (NIH).

#### Immunofluorescence analysis

Tumors embedded in OCT, a cryo embedding matrix, were cut as 10 μm thick section and fixed in ice cold methanol. Paraffin-embedded tumor sections were treated for antigen retrieval with sodium citrate buffer (Thermo Scientific Fisher). All tumor sections were blocked in blocking buffer containing PBS (Sigma), 0.3 % Triton X-100 (Sigma), 10 % fetal bovine serum (Gibco) and 1 % Bovine Serum Albumin (Sigma), and immunostained with the appropriate antibodies: anti CD68 (Dako), VEGFA (SantaCruz), CD31 (BD Biosciences), Podoplanin (Thermo Fischer Scientific), Lyve-1 (Novus Biologicals), NG2 (Merck Millipore), α-SMA (Sigma-Aldrich), F4/80 (Bio-Rad), VEGF-C (Thermo Fischer Scientific), VEGFR3 (R&D Systems), CD31 (DAKO), FITC-Lectin (Bio-Rad) and MENA/ENAH (Sigma). The appropriate secondary antibodies were conjugated with AlexaFluor 488, AlexaFluor 546 or AlexaFluor 647 fluorochromes (Life Technologies). Cell nuclei were labelled with DAPI (Sigma Aldrich). 10 independent fields from each section were analyzed by using LSM 710 Zeiss confocal microscope and quantified either using ImageJ or QuPath softwares.

#### Immunohistochemical analysis

##### Tumor hypoxia and vessel functionality

Tumor hypoxia was assessed by intraperitoneal injection of 60 mg/kg pimonidazole hydrochloride (Hypoxiprobe) into tumor-bearing mice. Mice were sacrificed 1 h after injection and tumors were harvested. To detect the formation of hypoxic adducts, OCT tumor sections were immunostained with Hypoxyprobe-1-Mab-1 (Hypoxiprobe kit, Chemicon) following manufacturer’s instructions. Vessel perfusion was evaluated by intravenous injection of 1 mg of FITC-conjugated Lectin (Thermo Fischer Scientific). Tumors were embedded in OCT material (VWR Chemicals) and snap-frozen in iso-pentane (VWR Chemicals) on dry ice.

### Quantification and statistical analysis

#### Image analysis

*Image J*-Image analysis was performed using FIJI/ImageJ. For quantifications related to hypoxia (**Figures 2H,3E,3O**), and dextran positive CD31 blood vessels (**Figure S3F**) (and counting for lymph node metastasis (**Figure 3J**) Regions of interest (ROI) were manually analyzed using the Polygon tool in ImageJ. The data were represented either as percent from total DAPI ROI (for the Hypoxia analysis) or presence of Dextran positive expression in the CD31^+^ blood vessels (for the Tie2 Dextran imaging). The final graphs were prepared and analyzed for statistics using graph pad prism software, version 10.

##### Qupath-analysis

Image analysis (**Figures 1D,1F,1H,1K,1M,1O,2B,2D,2E,2G,2K,2M,3B,3D,3F,3N,3R, S1C,S1E,S2B**) was performed by using QuPath 0.4.3 ^48^. Vessel segmentation was performed by using a pixel classifier based on the CD31 channel with a threshold allowing detection of both low and high CD31 signal to segment both lymphatic and blood vessels. For vessel classification, different object classifiers were created for each marker, allowing automatic classification of all vessel objects. To determine the marker coverage of the vessel area, pixel classifiers were created. For the analysis of macrophages, the cell segmentation tool was used together with object classifiers for each marker. Unique thresholds were used for each marker and staining. The final graphs were prepared and analyzed for statistics using graph pad prism software, version 10.

#### Financial support

This study was supported by the Swedish Cancer Society (2016/825, 2019/0281 and 2022/2384), Swedish Scientific Council (VR, 2018-02915 and 2021-01149), Radiumhemmets research funds 2016, 2019, 2021) to CR; Swedish Cancer Society and the Wallenberg foundation to JB; TW was supported by KI-PhD (2014-2018) foundation. JH was supported by grants from Radiumhemmets research funds, Swedish Breast Cancer Association, Swedish Society for Medical Research. MLS was supported by Fondazione Regionale per la Ricerca Biomedica (FRRB) 1751658-2020 and CARIPLO 1834-2019.

All the authors declare no conflict of interest regarding the work that is being submitted.

## Supplementary information

**Supplemental Figure 1.**
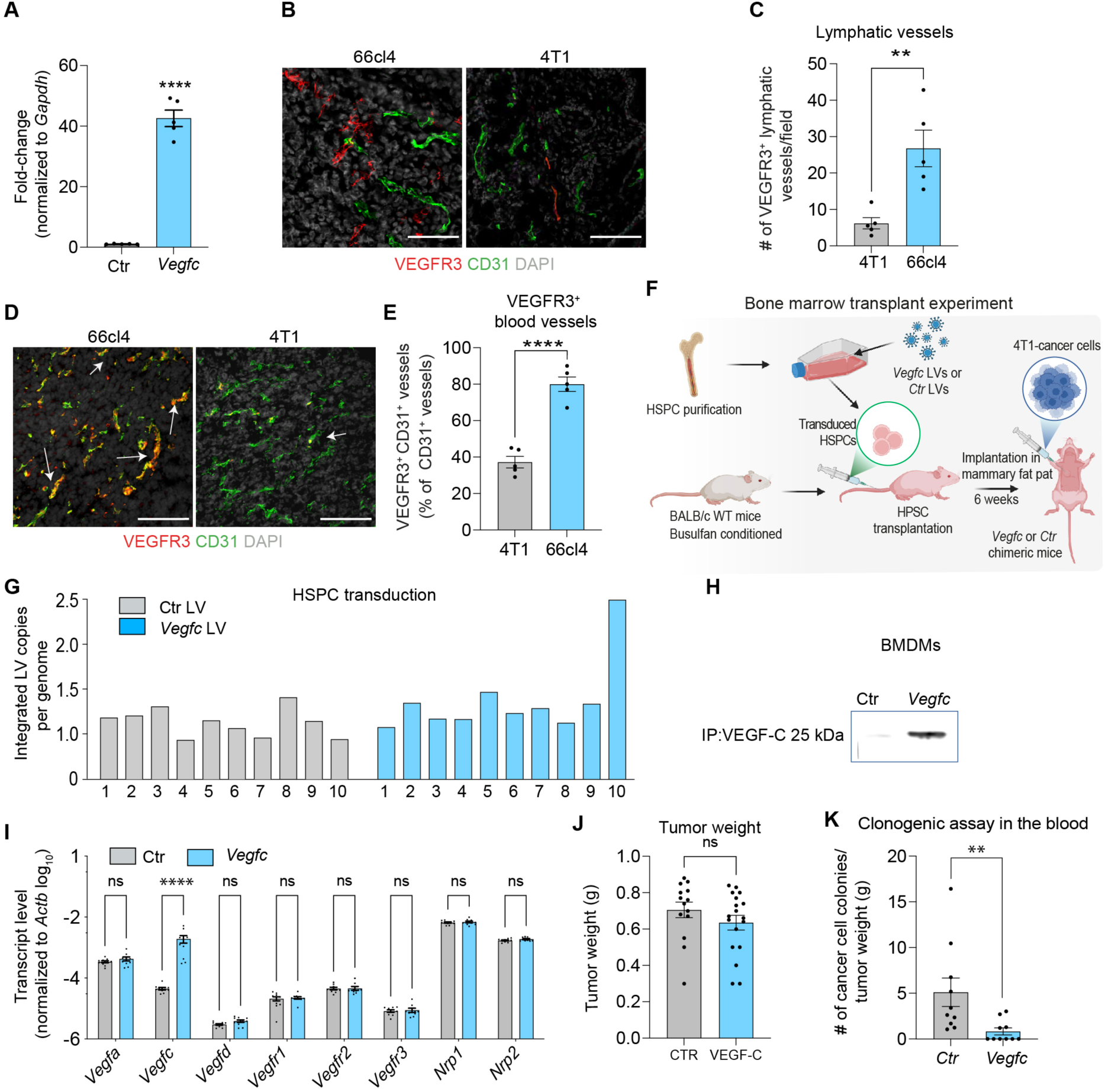
Perivascular VEGF-C expressing TAMs. Related to Figure 1. (A), RT-PCR analysis of *Vegfc* gene expression in BMDMs transduced with Ctr and *Vegfc* LVs. Statistical analyses were performed by Students t-test, *indicates significance; *p<0.05, n=5. All data are presented as the mean + SEM. (B-E), 66cl4 and 4T1 tumor sections were immunoassayed for CD31 (green) and VEGFR3 (red) in (B and D). Number of VEGFR3^+^CD31^low^ vessels were quantified out of vessels expressing VEGFR3 in (C) and area of VEGFR3^+^ expression of CD31^high^ vessels are depicted in (E). Statistical analyses were performed by Students t-test, *indicates significance; **p<0.01; ***p<0.001, n=5. All data are presented as the mean + SEM. (F-K), 4T1 tumor cells were injected into the mammary fat pad of syngeneic chimeric mice harboring haemopoietic cells engineered to express *Vegfc* or only the lentiviral backbone (Ctr). (F), Schematics describing the experimental procedure. (G), Genomic DNA was analyzed by qPCR for lentiviral integration of *Vegfc* or Ctr in bone marrow stem cells from CTR and VEGF-C chimeric mice. Histogram depicts lentiviral copies per genome. n=10. (H), VEGF-C protein expression in BMDMs derived from CTR and VEGF-C chimeric mice was analyzed by western blot. (I), RT-PCR analysis of gene expression for the VEGF family and its co-receptors in BMDMs attained from *Ctr* and *Vegfc*-chimeric mice. Statistical analyses were performed by Students t-test, *indicates significance; ****p<0.0001 not significant is indicated by ns, n=10. All data are presented as the mean + SEM. (J), Histogram depicts Ctr and Vegfc tumor weight (g). Statistical analyses were performed by Students t-test, not significant is indicated by ns, n=14-18. All data are presented as the mean + SEM. (K), The graph displays the metastatic index of blood tumor cell colonies (number of metastatic colonies in blood/g of primary tumor weight) of Ctr or *Vegfc*-chimeric tumor bearing mice. Students t-test, *indicates significance; **p<0.01, n=10. All data are presented as the mean + SEM.

**Supplemental Figure 2.**
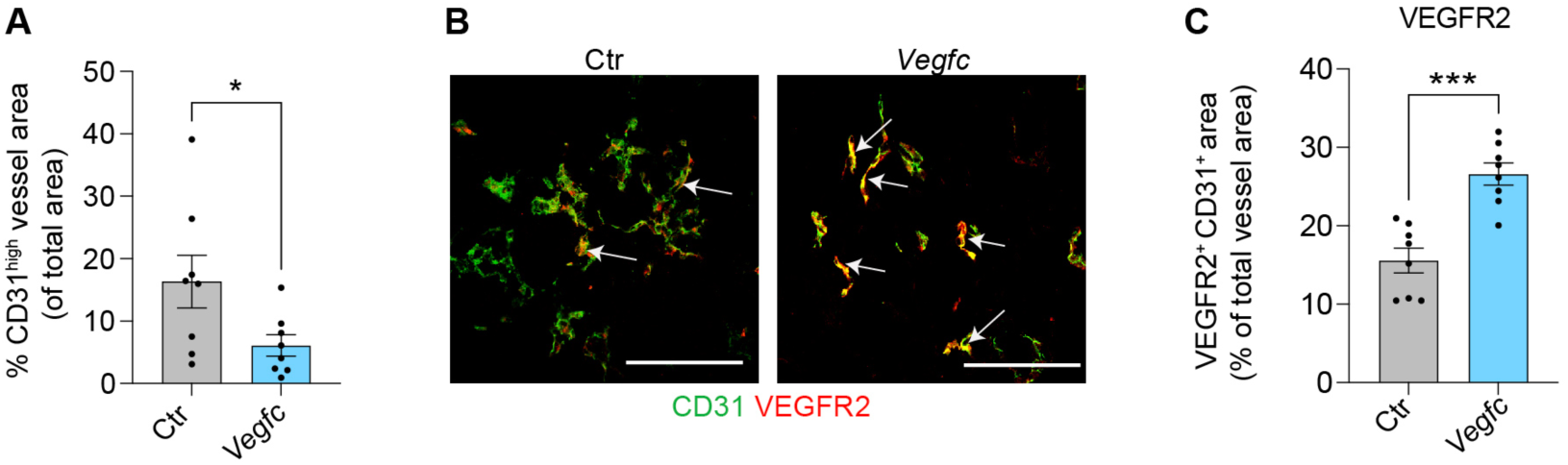
VEGFR2 expression in tumor endothelium. Related to Figure 2. 4T1 tumor cells were injected into the mammary fat pad of syngeneic chimeric mice harboring haemopoietic cells engineered to express *Vegfc* or Ctr. (A) Graph depicts morphometric analysis of CD31^high^ vessel area in Ctr and *Vegfc*-tumors (B-C) Ctr and *Vegfc*-4T1 tumor sections were immunoassayed for CD31 (green) and VEGFR2 (red) in (B). Area of VEGFR2^+^ expression of CD31^+^ vessels are depicted in (C). Statistical analyses were performed by Students t-test, *indicates significance;; ***p<0.001, n=8. All data are presented as the mean + SEM.

**Supplemental Figure 3.**
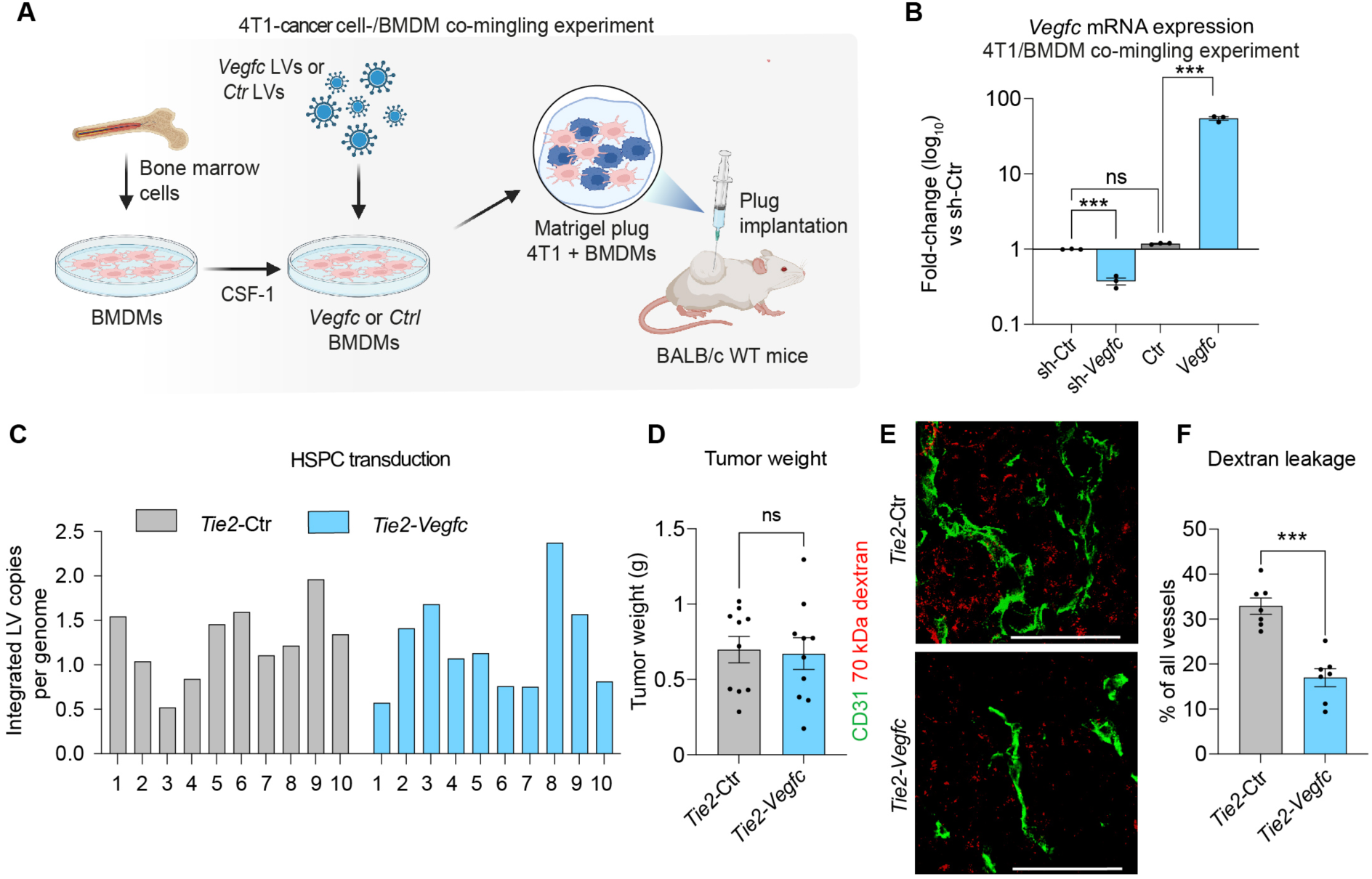
Targeted modulation of VEGF-C expression in TAMs rewires the metastatic destiny of 4T1 cells. Related to Figure 3. (A), Schematic figure of 4T1 cancer cells were co-mingled in matrigel with BMDMs transduced with Ctr or *Vegfc*-LVs and injected subcutaneously into syngeneic mice. (B), RT-PCR analysis of gene expression for the VEGF-C in BMDMs attained transduced with Ctr, *Vegfc*, sh-Ctr or sh-*Vegfc* LVs. Statistical analyses were performed by One-way ANOVA with paired analyses, *indicates significance; ****p<0.0001 not significant is indicated by ns, n=10. All data are presented as the mean + SEM. C-F, 4T1 cancer cells were injected into the mammary fat pad of syngeneic chimeric mice harboring haemopoietic cells transduced with *Tie2-*Ctr or *Tie2-Vegfc* lentiviral vectors in (C), Genomic DNA was analyzed by qPCR for lentiviral integration of *Vegfc* in bone marrow stem cells from CTR and VEGF-C chimeric mice. Histogram depicts lentiviral copies per genome. n=10. (D), Graph shows tumor weight derived from *Tie2*-Ctr and *Tie2-Vegfc* chimeric mice. Statistical analyses were performed by Students t-test, not significant is indicated by ns, n=10. All data are presented as the mean + SEM. (E-F), 4T1 tumor bearing *Tie2*-Ctr and *Tie2-Vegfc* chimeric mice were injected with 70 kDa Dextran intraperitoneally before sacrificing. Representative images show immunostaining for CD31 (red), Streptavidin-conjugated AF555 (red) Graph shows quantification from % of Dextran leakage from CD31+ blood vessel n=7 mice/group. Statistical analyses were performed by Students t-test, stars indicate significance ***p<0.001, All data values are shown as the mean ± SEM. Representative images are shown. At least 5-6 OFs are visualized per tumor section and representative images are shown. Bars represent 50 μm.

**Supplemental Figure 4.**
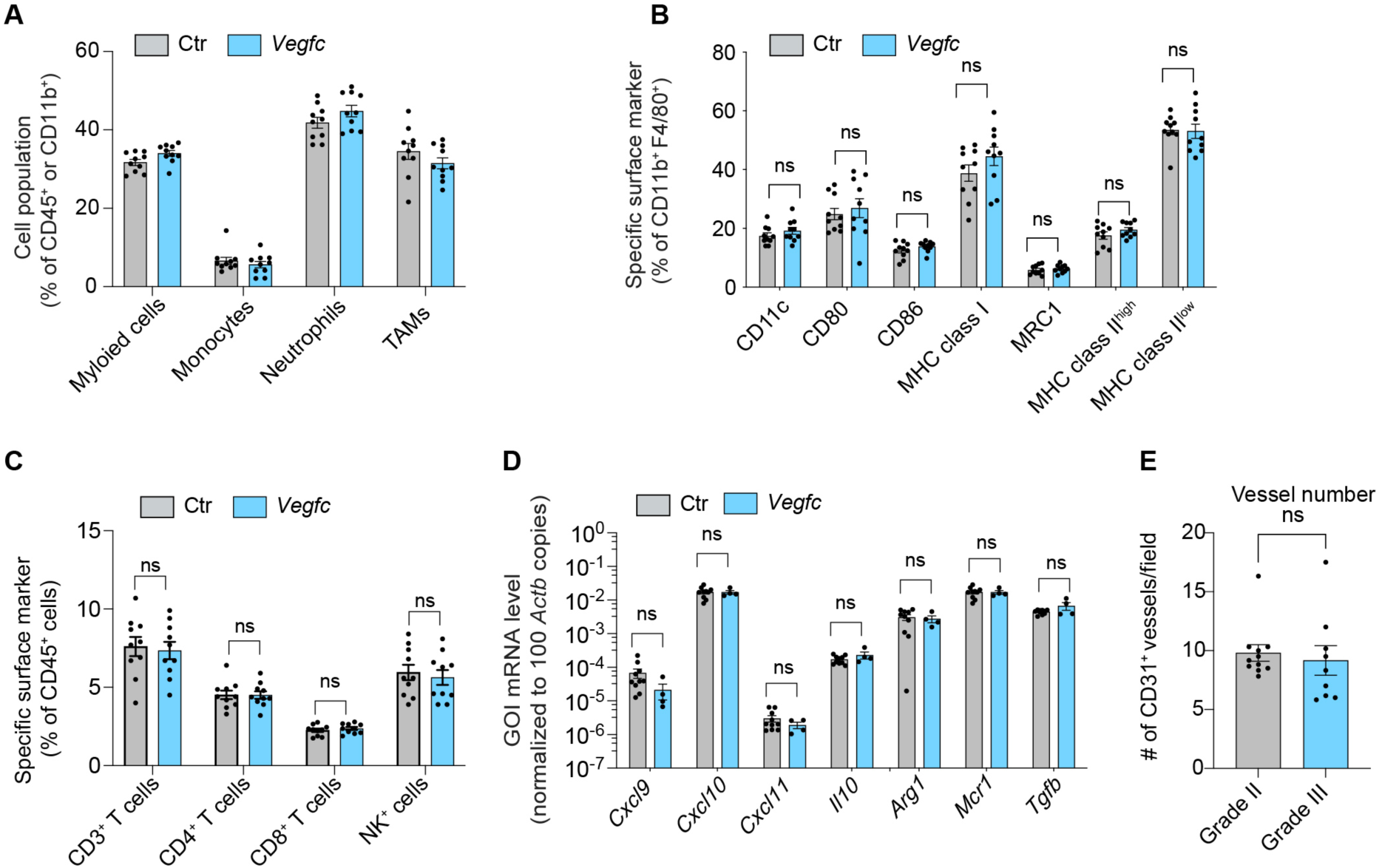
TAM-derived VEGF-C does not the affect myeloid and lymphoid immune landscape. Related to Figure 4. 4T1 tumor cells were injected into the mammary fat pad of syngeneic chimeric mice harboring haemopoietic cells engineered to express *Vegfc* or Ctr. A-C, Graphs show flow cytometry analysis of 4T1-CTR and 4T1-VEGF-C tumors for myeloid cells (out of CD45), TAMs, neutrophils, monocytes in (A) TAMs phenotypes in (B), T cells and NK cells in (C) D, Graph displays quantitative RT-PCR analysis of gene expression levels of BMDMs derived from Ctr- or *Vegfc*-mice. Gene expression was normalized to β-actin levels. (4E) The graph depicts morphometric analysis of BC specimens of grades II and III immunostained for CD31, n>4-16. Statistical analysis by students t-test was not significant (NS).

